# Systematic exploration of unsupervised methods for mapping behavior

**DOI:** 10.1101/051300

**Authors:** Jeremy G. Todd, Jamey S. Kain, Benjamin L. de Bivort

## Abstract

To fully understand the mechanisms giving rise to behavior, we need to be able to precisely measure it. When coupled with large behavioral data sets, unsupervised clustering methods offer the potential of unbiased mapping of behavioral spaces. However, unsupervised techniques to map behavioral spaces are in their infancy, and there have been few systematic considerations of all the methodological options. We compared the performance of seven distinct mapping methods in clustering a data set consisting of the x-and y-positions of the six legs of individual flies. Legs were automatically tracked by small pieces of fluorescent dye, while the fly was tethered and walking on an air-suspended ball. We find that there is considerable variation in the performance of these mapping methods, and that better performance is attained when clustering is done in higher dimensional spaces (which are otherwise less preferable because they are hard to visualize). High dimensionality means that some algorithms, including the non-parametric watershed cluster assignment algorithm, cannot be used. We developed an alternative watershed algorithm which can be used in high-dimensional spaces when the probability density estimate can be computed directly. With these tools in hand, we examined the behavioral space of fly leg postural dynamics and locomotion. We find a striking division of behavior into modes involving the fore legs and modes involving the hind legs, with few direct transitions between them. By computing behavioral clusters using the data from all flies simultaneously, we show that this division appears to be common to all flies. We also identify individual-to-individual differences in behavior and behavioral transitions. Lastly, we suggest a computational pipeline that can achieve satisfactory levels of performance without the taxing computational demands of a systematic combinatorial approach.

**Abbreviations:** GMM: Gaussian mixture model; PCA: principal components analysis; SW: sparse watershed; t-SNE: t-distributed stochastic neighbor embedding

## INTRODUCTION

Understanding how nervous systems integrate information from the environment, past experience and internal states to produce useful behaviors is a key goal of behavioral neuroscience. Rapid progress is being made toward this goal, especially using animal models with relatively simple brains. Large quantities of neural activity data have been acquired simultaneously with behavioral data in larval zebrafish *Danio rerio* (Dunn *et al* 2016), the nematode *Caenorhabditis elegans* (Nguyen *et al* 2016, Venkatachalam *et al* 2016), and the fruit fly *Drosophila melanogaster* (Lemon *et al* 2015, Harris *et al* 2015, Seelig and Jayaraman 2015). These organisms are also the focus of past (White *et al* 1986) or ongoing (Takemura *et al* 2013) connectomic efforts to map the synapse-level connectivity between all neurons in the brain. While analyses of neural activity and connectivity have been quantitative and richly multidimensional since their inception, the quantification of behavior is comparatively depauperate. For a full accounting of the neural basis of behavior, rich, detailed and unbiased descriptions are needed.

Manual annotation simply cannot produce data sets large enough to be analyzed in conjunction with gigabytes to petabytes of neural activity and connectomic data. Approaches to automated behavioral classification can be roughly sorted into *supervised methods*, in which an investigator-labeled training set is used to train classifiers that can then assign those labels to behavioral instances not included in the training set. By contrast, *unsupervised methods* seek to describe axes of variation or clusters of behavioral patterns with as few a priori assumptions as possible. The principal limitations of supervised methods are manually building up a large enough training set to achieve high classifier performance and not being able to detect behavioral patterns beyond the categories stipulated *a priori*. On the other hand, it is hard to evaluate the performance of unsupervised methods because to the extent they impose very few assumptions, they have no internal benchmarks of success. Instead, overall success is normally assessed by manual, qualitative inspection of the clustered behavioral instances.

Unsupervised methods have been applied successfully to diverse questions in neuroscience. These include clustering modules of genes by their association with behavioral states in social insects (Chandrasekaran *et al* 2011), the identification of types of subtypes of retinal cells based on molecular diversity (Marc and Jones 2002, Macosko *et al* 2015) and cortical cells by expression profiles (Tasic *et al* 2016), sorting of action potential recordings (Quiroga *et al* 2004), and the sorting of neurons by the three-dimensional morphology of their axonal and dendritic processes (Costa *et al* 2014). Particular progress has been made in the analysis of the behavior of *C. elegans*, in part because its posture can be faithfully compressed into a low four-dimensional representation, termed “eigenworms” (Stephens *et al* 2008). This discovery led to rapid progress in the unsupervised classification of *C. elegans* postural trajectories and their defects in mutant animals (Yemini *et al* 2013, Brown *et al* 2013, Schwarz *et al* 2015), as well as the biophysical basis of postural dynamics (Stephens *et al* 2011).

In *Drosophila melanogaster*, a long ethological history (e.g. Waddington *et al* 1954, Hirsch and Erlenmeyer-Kimling 1961, Chen *et al* 2002) predates methods for automated behavioral analysis. Early efforts to automate fly behavioral analysis naturally sought to facilitate what would otherwise be manual efforts, in particular the classification of video data. These approaches produced widely used tools for tracking adult flies freely behaving in an arena at relatively high density (Branson *et al* 2009, Dankert *et al* 2009) and larvae at lower densities for exhibiting non-social behaviors (Luo *et al* 2010). This trend has culminated in the release of versatile turn-key tools for quickly collecting training set data and automatically training classifiers (Kabra *et al* 2013). The sophistication of supervised methods continues to increase with the added estimation of underlying behavioral transition models (Saeedi *et al* 2016).

Applications of unsupervised methods to map fly behavior are rare by comparison. Geurten *et al* (2010) used cluster analysis to identify discrete components of hoverfly flight trajectories. In 2014, two impressive papers employed unsupervised behavioral clustering in *Drosophila*. Berman et al., reported the first unsupervised mapping of adult *Drosophila* behavior from video data using probability density estimation to identify modes in time-frequency transformed data, thereby identifying stereotyped postural dynamics. Vogelstein *et al* (2014) used unsupervised structure learning to infer a hierarchical organization of larval behaviors based on eight time varying measures of posture and motion. Most recently an “eigenlarva” analysis revealed that many behavioral events defy easy assignment to discrete clusters, suggesting that, at least in larvae, behavior may vary rather continuously (Szigeti *et al* 2015).

Here, our goals are two-fold. First we want to begin the systematic comparison of alternative approaches for unsupervised clustering. Despite the lack of a built-in performance metric (e.g. % correct classification), we believe that head-to-head comparison of unsupervised methods is possible using measurable qualities we expect from successful behavioral clusterings. Second, we want to apply the method that appears best to a data set of adult *Drosophila* leg-positions and motion vectors. Measurements come from both wild type flies and animals bearing mutations in the gene *nanchung*, which encodes a TRP channel that mediates proprioceptive sensation (Gong *et al* 2004). Thus, our aim is to find the best method to produce a model-free map of the space of postures and postural dynamics exhibited by these animals.

We recognize that many methods for unsupervised clustering of behavioral data (and presumably time-series data in general) share a common architecture. 1) First, *machine vision* is performed to capture behavior in several dimensions. 2) This data then undergoes *pre-processing* and may be dimensionally expanded using time frequency analysis to capture postural dynamics (e.g. Berman *et al* 2014 and Wiltschko *et al* 2015). 3) *Dimensionality reduction* is employed to facilitate subsequent computational steps. 4) Lastly, a *cluster assignment* algorithm is used to assign individual data frames to a discrete list of behavioral modes. Some clustering algorithms require an intermediate *density estimation* step that approximates the underlying probability distribution that gives rise to the data, e.g. the watershed algorithm (Meyer 1994) and Gaussian mixture modeling (GMM) posterior probability estimation (McLachlan and Peel 2000). Other clustering algorithms do not require density estimation (e.g. k-means clustering (Lloyd 1982)).

Importantly, approaches with this architecture are modular. Alternative algorithms can be used at each stage interchangeably. Principal components analysis (PCA; Pearson 1901) and t-distributed stochastic neighbor embedding (t-SNE; van der Maaten and Hinton 2008) both implement dimensionality reduction with respective advantages and disadvantages (van der Maaten *et al* 2009). Both of these algorithms (and many others that will not be considered further, such as multidimensional scaling (Torgerson 1952), isomap (Tenenbaum *et al* 2000)) can serve the role of reducing dimensionality. These alternatives can be combinatorially mixed with the alternative algorithms for cluster assignment to yield a large number of possible unsupervised clustering methods.

Moreover, the degree of dimensionality reduction can be tuned continuously, though not arbitrarily, as some downstream steps may be computationally intractable without sufficient dimensionality reduction. For example, the watershed algorithm examines the probability densities associated with every point in a space, down to some level of granularity. If each dimension is resolved into 500 bins, as would make for a nice looking image, then the number of points that must be examined is 500^*d*^ where *d* is the dimensionality of the space. Even with such dimensionality constraints, it is plausible that the degree of dimension reduction prior to clustering matters for the performance of the overall method, perhaps critically. There have been few systematic efforts to examine an assortment of methods. The work of Wiltschko *et al* (2015) is an exception, as they consider several methods to model the temporal evolution of mouse behavior.

Here we examine two alternative methods for dimension reduction, PCA and t-SNE. These algorithms represent opposite ends of the spectrum ranging from projection/ embedding algorithms that prioritize the preservation of global structure (PCA) to those that prioritize the preservation of local structure (t-SNE). In a combinatorial fashion, we consider two different cluster assignment methods: watershed and GMM posterior probabilities. These two algorithms each require a density estimation step, and we use 2D Gaussian blurring and Gaussian mixture modeling for them respectively.

Objectively comparing unsupervised clustering methods is philosophically challenging (Jain *et al* 1999), as unsupervised methods attempt to make useful clusters with as few as possible *a priori* assumptions. Identifying an unsupervised method as successful because it produces any particular quality in its output implicitly constrains the types of classifications that can be made, and pushes the method in the direction of supervised clustering/classification. Nevertheless, we claim that it is reasonable to posit very general desirable properties of a clustering output. For example, a “better” clustering method will produce fewer bouts of behavior that are impossibly short, i.e. faster than the animal can implement physically. Perhaps a better clustering method would also allow improved prediction of upcoming behavioral states given current states. In this way, we posited several qualities of good clustering methods, developed numerical metrics to capture them, and applied them to compare the various unsupervised clustering methods possible using the separate steps described above.

With these metrics as guides, we systematically examined seven unsupervised clustering methods. One of these was clearly the best (PCA reduction to 20 dimensions followed by GMM, with the addition of watershed-inspired cluster consolidation), by our metrics. With it we mapped the space of posture dynamics in a data set that measured the x-and y-position of flies’ legs as they walk, tethered above a floating ball.

## METHODS

### Data and code

All code and data used in this project will be available for download at *http://lab.debivort.org/unsupervised-methods-for-mapping-behavior*. The raw data will be also available at *http://zenodo.org/URLREFREF*. The code is also available at *https://github.com/de-Bivort-Lab/behavior-mapping*.

### Fly husbandry and experiments

Female Canton-S (wild type) and *nanchung*^*36a*^ (*nan*; Bloomington stock #24902) flies were reared on standard CalTech recipe cornmeal-based growth medium, in 23°C incubators at approximately 40% humidity and on 12h-12h light-dark cycles. 2-6 day old animals were marked with dye as described elsewhere (Kain *et al* 2013), and recorded on the leg-tracker for 2h at room temperature and approximately 15% humidity, under white LED lighting. Digital video was acquired for the first 5 minutes of each wild type experiment using a filter which blocked the HeNe laser emission used to excite the dye. Visible LED illumination passing this filter appeared purple and emitted IR dye fluorescence appeared orange. Flies were identified by sequential numbers corresponding to experiments. Recordings pertinent to this study began with experiment 37. With all flies, we attempted to recover the animal (unglue it from its tether) for repeated longitudinal testing across days. This yielded 2 sequential recordings, separated by ∼24h, for 6 wild type flies and 3 sequential recordings, separated by ∼24 and ∼48h for two wild type flies. Four additional wild type animals yielded a single recording each. Five *nan* flies were tested in total, none of which yielded sequential recordings.

### Data pre-processing and error correction

The raw data from the instrument (Kain *et al* 2013) was preprocessed before being subjected to analysis. First we used a 3-frame median filter to reject single-frame noise in the data, then we resampled the variable-rate instrument data to 100 Hz. To detect errors longer than a single frame we computed σ, the standard deviation over the entire experiment. Frame-to-frame changes greater than 5σ were flagged as errors until the leg returned to within 5σ of its pre-error position or within 1σ of the overall median position.

### Gaussian mixture models

All Gaussian mixture models in this paper were fit using MATLAB’s built-in expectation-maximization algorithm with fully independent and unconstrained covariance matrices. To estimate the scale of Gaussian mixture model components we examined the eigenvalues of the corresponding covariance matrix. The eigenvalues correspond to the standard deviation of the distribution along its principal axes (Banfield and Raftery 1993), so we multiply them together to estimate the relative scale of each mixture component. When co-fitting data from multiple fly experiments with a Gaussian mixture model we built a training set with 2 million frames by decimating the frames from each trial so that an equal number of frames were taken from every trial. A PCA transform was computed using the training set and applied to the full data set, and the resulting mixture model was used to classify the full data set in PCA space.

### Computing

All of our analysis was done using built-in functions and custom code run in MATLAB R2015b-2016a. Computationally intensive portions of mapping methods (t-SNE, GMM in high dimensions, and sparse watershed) were run on the Odyssey cluster (FAS Division of Science, Research Computing Group, Harvard University). The rest of the processing and figure preparation was done on a MacBook Pro with 16 GB of RAM.

### Statistics and clustering

Significant differences between mapping method metric values across flies were determined by pairwise Wilcoxon signed-rank tests, corrected for 90 comparisons using the formula *p*’=1-(1-*p*)^90^. For comparisons of PCA_20_-GMM-SW and PCA_20_-GMM, see below, metric values, correction was applied for 6 comparisons. Statistical significance of metric value differences between wild type and *nan* mutants were calculated using Wilcoxon–Mann–Whitney tests corrected for 36 comparisons (no such differences were significant). Metric mean Coefficients of Variation were calculated across 5 independently random seeded replicates of each fly x mapping method x metric, and then averaged across flies. Wilcoxon–Mann–Whitney tests were used to compare measures within a fly across trials with measures between random pairs of flies, as well as differences in cluster abundance between genotypes, with the latter corrected for 41 comparisons. Matrix row and column clustering was determined using the *linkage* command in MATLAB, using the ‘spearman’ method, which minimizes distortion from very large input dimensions.

## RESULTS

### Unsupervised clustering methods

We refer to our unsupervised clustering methods collectively as *mapping methods*, and they all share a common structure. They begin with a data preparation step (common to all mapping methods), followed by dimensionality reduction, density estimation and finally cluster assignment. We name each of them with an acronym for 1) the dimensionality reduction method, 2) a subscript indicating the number of resulting dimensions, and 3) the cluster assignment algorithm. For example the method in Berman *et al* (2014) is *tSNE2-watershed* mapping as it uses t-SNE to reduce the data to two dimensions and a watershed transform for clustering (with an intermediate density estimation step done by 2D Gaussian blurring which we omit from the acronym). The six mapping methods that are the focus of this analysis are schematized in Figure 1A. In addition, as a baseline comparison class, we also implemented a *random* mapping method in which frames are independently assigned at random to one of *k* clusters.

**Figure 1.**
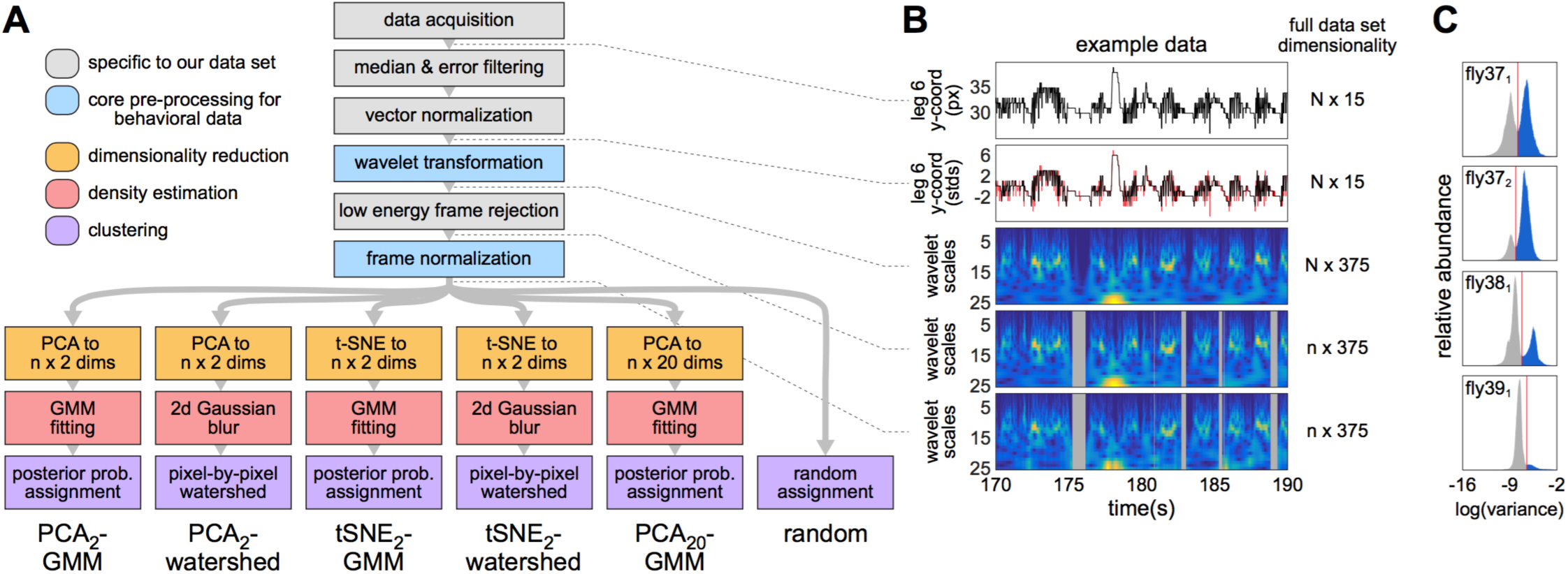
Unsupervised methods for mapping Drosophila behavioral space –. **A**) Flow chart diagrams of the 6 combinatorial mapping methods considered herein. Box colors indicate stages common to all methods. **B**) Example data from fly experiment 37_1_ illustrating each of the successive stages of pre-processing prior to dimensionality reduction. Dotted lines connect the example data to the point in the method where the data existed in that form. Red line in the second panel indicates the data prior to filtering and normalization (i.e. the data in the first panel). Time-frequency heatmaps in panels 3-5 are ordered from high frequencies at the top to low frequencies at the bottom. Grey blocks in the last two panels indicate low-variance frames that were rejected from further analysis. **C**) Histograms of frame-by-frame variance in wavelet energies (i.e. the variance of the 375 dimensions at each frame in the middle panel of B). Red line indicates the threshold below which frames were rejected (grey). The first 5 histograms from our data set are shown as a representative sample.

### Data preparation

Our data consists of a 15-dimensional 100 Hz time series: the x-and y-coordinates of each of the six legs, and the three rotational velocities of the trackball as published by Kain *et al* (2013) and five recordings from *nan* flies which are new to this publication.

The first step in all of our mapping methods is a wavelet transform which might be included in most analyses of behavior. The wavelet domain is a useful representation of our data for reasons given by Berman *et al* (2014): it describes dynamics over several timescales simultaneously, and by taking the magnitude of the complex wavelet coefficients it phase-aligns periodic behaviors (Figure 1B). We used the wavelet transform used by Berman *et al* (2014) with 25 wavelet scales, spaced logarithmically between 1 and 50 Hz (the Nyquist limit). This yielded a 375-dimensional (15*25=375) time-series (see Supplemental Movie M1 for an example of time domain and wavelet data, along with subsequent steps in our mapping methods).

As described by Berman *et al* (2014), the magnitude of wavelet coefficients can vary due to the wavelet’s analysis window. To compensate for this we normalized each frame by the sum of its wavelet magnitudes. Thus we analyzed the relative energy at each wavelet scale (roughly equivalent to a frequency band) in our data (Figure 1B). The 375-dimensional frame-normalized wavelet space captures how the fly’s posture changes over time, thus we refer to it as *postural dynamics space*.

Our data is mean-centered, and when the animal is at rest it tends to a zero value. So rest frames contain only noise energy. Frame normalization amplifies this low-level noise energy, generating a wide range of random high-energy behaviors. To avoid this, we noted that the distribution of frame variances is bimodal (Figure 1C), corresponding to rest frames and frames with activity. We thus classified frames as either low-variance or high-variance based on these modes, and we subjected only high-variance frames to frame normalization and subsequent analysis. For the purposes of cluster assignment all low-variance frames were assigned to a single cluster. This approach also had the benefit of reducing computational requirements, since roughly half of the frames in our data set were classified as rest frames.

### Dimensionality reduction

Berman *et al* (2014) (and several of our mapping methods) reduce data from a high-dimensional space to a two-dimensional *t-SNE space*. t-SNE (van der Maaten and Hinton 2008) is a nonlinear embedding that attempts to preserve the local structure of the high-dimensional space, as opposed to the global structure which is preserved by a linear embedding such as PCA. This means that two points which are close in the high-dimensional space should have their proximity preserved in the t-SNE space, while two points which are distant in the high-dimensional space may or may not end up far apart in the t-SNE space. For clustering applications this property is desirable, since clusters are defined by local structure, and moving clusters around (i.e. changing global structure) in the t-SNE space ought not to have a significant effect on cluster assignments. Once data points are embedded in the t-SNE space, they can be visualized, which is a very convenient property of the mapping methods that reduce data to two dimensions.

Our alternative technique for dimensionality reduction is principal component analysis (PCA). PCA selects an orthogonal set of basis vectors which maximize the amount of variance captured when the data is projected onto this basis. In order to determine how many principal components to keep we used the shuffling procedure described by Berman *et al* (2014) to quantify sampling error. The procedure indicated that between 10 and 20 (depending on the fly experiment) principal components contained variance above sampling error and should be kept, explaining 60-70% of the variance in the data. We decided to keep 20 principal components for all trials to facilitate comparisons between trials and to make sure all relevant variance was captured. We refer to the resulting 20-dimensional space containing PCA scores as *compressed postural dynamics space*. For comparison with t-SNE we also use PCA to reduce our data to two dimensions in several of our mapping methods.

### Density estimation

Once we have a space with suitable dimension, the next step is to estimate the density of our data in this space. This can be viewed as a statistical procedure converting our finite set of data samples into a continuous probability density function from which animals sample their behavior.

One technique for density estimation is to fit our data points with a Gaussian mixture model (GMM). A Gaussian mixture model assumes data points are drawn from a probability distribution formed by summing mixture components. Each mixture component is a multivariate Gaussian distribution with dimension matching our space (e.g. 20 dimensions in the case of our compressed postural dynamics space) with a weight (a scalar), a multivariate mean (a 20-dimensional vector) and a covariance matrix (a 20x20 matrix). For the purposes of mapping method comparison we chose the number of mixture components (the key fitting parameter of a GMM, usually denoted *k*) to match the number of clusters found in other methods. The fitting procedure attempts to find the model which maximizes the likelihood of our data given the GMM.

The density estimation technique used by Berman *et al* (2014) applies a 2D Gaussian filter to the points in t-SNE space, yielding a 501 x 501 pixel *density map*. The width of this Gaussian filter can be tuned to produce a wide range of cluster counts, but we use the default width used by Berman *et al*.

### Cluster assignment

When density estimation is done by fitting a GMM, each data point has a set of posterior probabilities, the respective probability of drawing that point from each mixture component. To assign a data point to a cluster we choose the mixture component with the maximum posterior probability (McLachlan and Peel 2000). This assignment is less clear if several mixture components have large posterior probabilities. As we will see below, this concern is not especially problematic in our data set.

The clustering method used by Berman *et al* (2014) is a watershed transform (Meyer 1994). It assigns two data points to the same cluster if they would both reach the same local maximum by ascending the local gradient. The watershed transform produces intuitive cluster assignments, but its memory usage scales with M, the number of bins along each dimension of the density map, and d, the number of dimensions, as M^d^. Thus it is feasible when our dimensionality reduction yields two dimensions, but it is not useful in our 20-dimensional compressed postural dynamics space.

### Metrics

It is difficult to compare results in unsupervised clustering problems because there is no general definition of a “correct” or “better” clustering (Jain *et al* 1999). Rather than attempt to create a single rule comparing clusterings, we assembled a set of quantitative metrics on clusterings. In some cases a high or low value for a metric is desirable for a good clustering. In other cases a metric is mostly of interest for trying to understand a given clustering. We evaluated each of the following metrics for each mapping method and each fly experiment (Figure 2).

**Figure 2.**
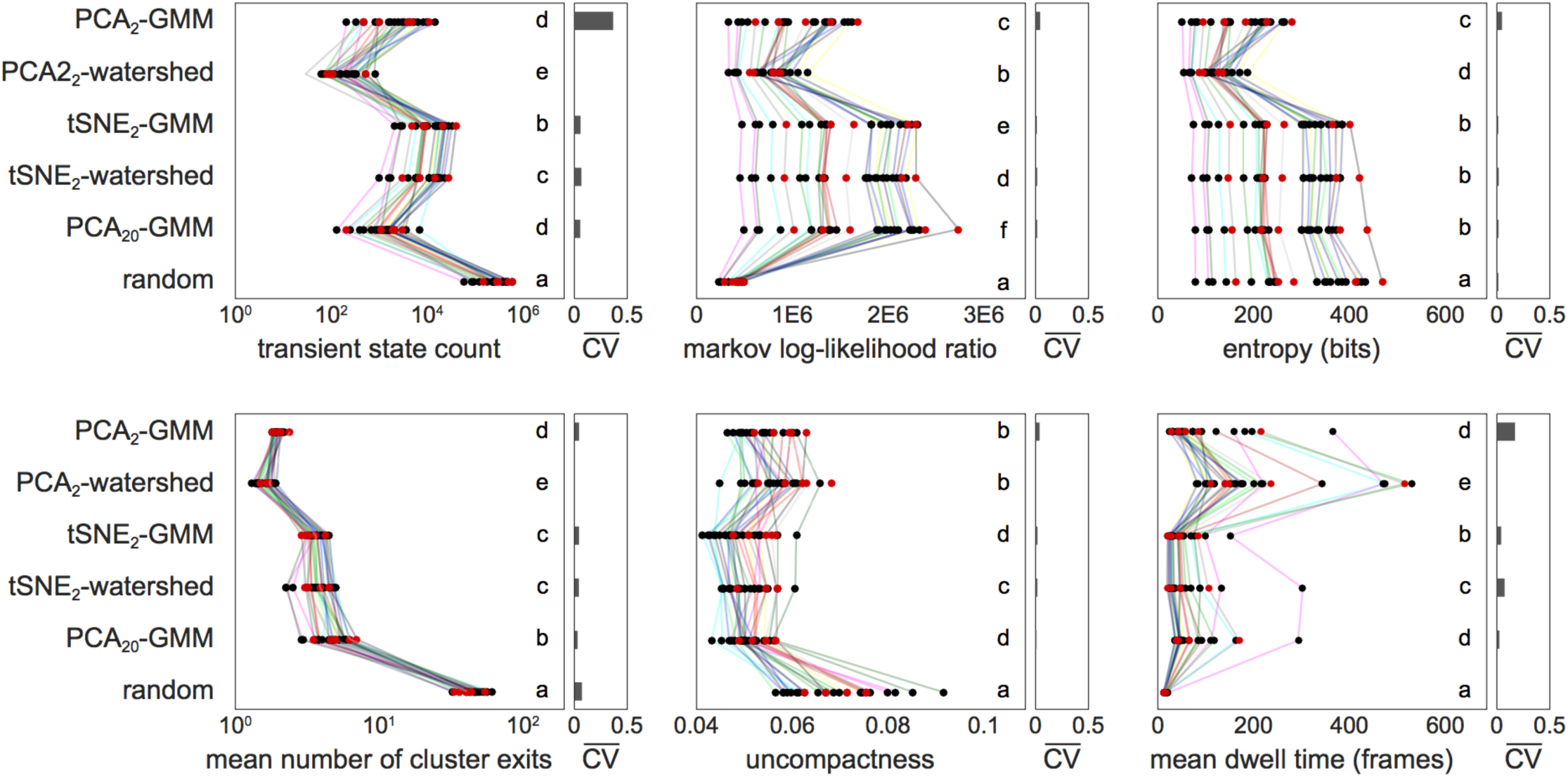
Comparison of metric values across mapping methods and flies –. Metric values are plotted for each fly experiment and each mapping method. See text for explanation. Red points indicate *nan* fly experiments. Lines connect corresponding points across mapping methods. Lines of the same color indicate multiple experiments done on the same fly across successive days. Letters indicate statistically significant differences, with groups that share the same letter being statistically indistinguishable. Significance determined using pairwise Wilcoxon signed-rank tests corrected for 90 comparisons (see Methods). Bar graphs at the right of each panel are the coefficients of variation, averaged across fly experiments, of five replicates of each mapping method computed with different random seeds.

#### Transient state count

We assume there is a lower bound on how long an animal can remain in a behavioral state. For the fruit fly we consider 20 ms to be a conservative lower bound on dwell time in any given state, as seen in previous supervised classifications (Kain *et al* 2013) and unsupervised clusterings (Berman *et al* 2014) of fly behavior, as well as other species (Wiltschko *et al* 2015). Our *transient state count* is the number of cluster dwell times of 20 ms (2 frames at 100 Hz) or less. Therefore, a lower transient state count corresponds to a clustering which is more biologically plausible.

#### Markov Log-Likelihood Ratio

One desirable property of a clustering is its predictive power. If we assume that fly behavior is Markovian, we can examine the likelihood of the cluster sequence under a Markov model. To measure this, we first fit a first-order Markov model to a given sequence of cluster assignments and compute the log-likelihood of the data under this model. The transition matrix is dominated by self-transitions (see the mean dwell time metric below), so to normalize for this we subtracted the log-likelihood of the data under a zeroth-order Markov model. The result is a log-likelihood ratio measuring the improvement in likelihood obtained by considering transitions between states. A larger value is desirable as it indicates a greater cluster predictability.

#### Entropy

Many of the other metrics will be affected by how evenly frames are distributed across clusters. For example, if a single cluster tends to dominate then one can expect fewer state transitions (this probably also represents a less useful clustering). To quantify this we counted the number of data points in each cluster and then normalized to form a probability mass function (PMF). This PMF indicates the probability that a data point drawn at random will be assigned to a particular cluster. We then measured the diversity of cluster sizes by computing the entropy of this PMF. Entropy is maximal when frames are evenly distributed among all clusters.

#### Mean number of cluster exits

If we form a Markov transition matrix from our cluster assignments, we can look at the average out-degree of clusters. A network with lower mean out-degree is probably desirable in that it facilitates the prediction of sequences of clusters. We measured this by sorting the transition probabilities (excluding the self-transition) from most probable to least probable and taking an average weighted by rank. The mean number of cluster exits is the mean of this weighted average across clusters. A lower mean number of cluster exits indicates a more predictable sequence of cluster transitions.

#### Uncompactness

A compact clustering is one with a relatively small distance between points within the same cluster. To compute *uncompactness* we first find the mean of each cluster’s data points. Then for each cluster we compute the mean distance to the cluster mean over all of the points in that cluster. Finally we take the mean across clusters as our uncompactness metric. Since uncompactness is greatly affected by cluster shape (hyperspherical clusters will tend to minimize uncompactness), it is not necessarily desirable to minimize this metric.

#### Mean dwell time

The *mean dwell time* is the mean number of frames the fly spends in each cluster before transitioning to the next cluster. The transient state count metric captures our assumption about a lower bound on dwell times, so the mean dwell time is foremost a descriptive metric.

#### Repeatability

The implementations of GMM and t-SNE chosen for this paper are non-deterministic, since both algorithms start from a random solution and converge to a more optimal one. To measure how repeatable the algorithms are, we ran each mapping method 5 times for each fly experiment and computed the coefficient of variation for each metric (Figure 2, bars at right in each panel). Repeatability is probably a desirable quality in a clustering.

In order to identify the mapping method most appropriate to our data set we examined these metrics in aggregate (Figure 3). We sorted the metrics from those that definitely measure desirable qualities of a mapping method to those that are more neutrally descriptive. We saw that PCA_20_-GMM yielded desirable metric values for nearly all metrics. The one exception was the mean number of cluster exits, which we considered to be a metric for which low values were probably desirable.

**Figure 3.**
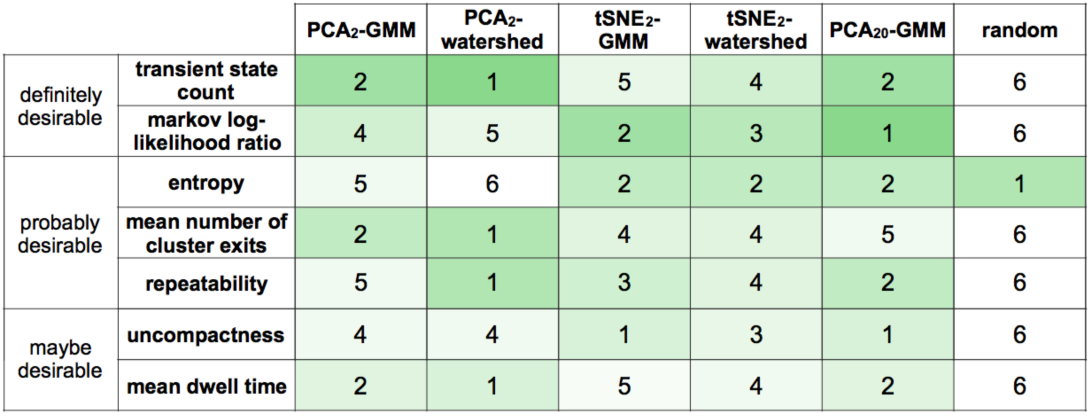
Summary of mapping method performance on all metrics –. Numbers indicate the rank of each method according to each metric. Methods are considered to be in different ranks if they are statistically significantly different. In evaluating repeatability we considered both the frame-by-frame stability of cluster assignments and the coefficient of variation measures in Figure 2. Lower rank numbers are preferable. Metrics are categorized by our confidence that the metric should be optimized for a method to be declared successful. Darker green shading indicates better ranks, with the darkest shades marking metrics where we have the greatest confidence that particular metric values are desirable.

### Cluster consolidation

Based on the above metrics we elected to focus on the PCA_20_-GMM mapping method. In order to gain intuition for the structure of the 20-dimensional compressed postural dynamics space, we first looked at the GMM covariance matrices to estimate the “volume” of each mixture component in this space (i.e. product of the eigenvalues of each component’s covariance matrix; Banfield and Raftery 1993). We found that cluster volumes are approximately log-normally distributed (Figure 4A), with a roughly exponential falloff in scale as a function of dimension (Figure 4B). That is, components are more linear or disk-shaped than high-dimensional balls. Since we use maximum posterior probability for cluster assignment, we also verified that the most likely mixture component tends to dominate the posterior probabilities for each frame of data, i.e., most frames are predominantly associated with just a single cluster (Figure 4C).

**Figure 4.**
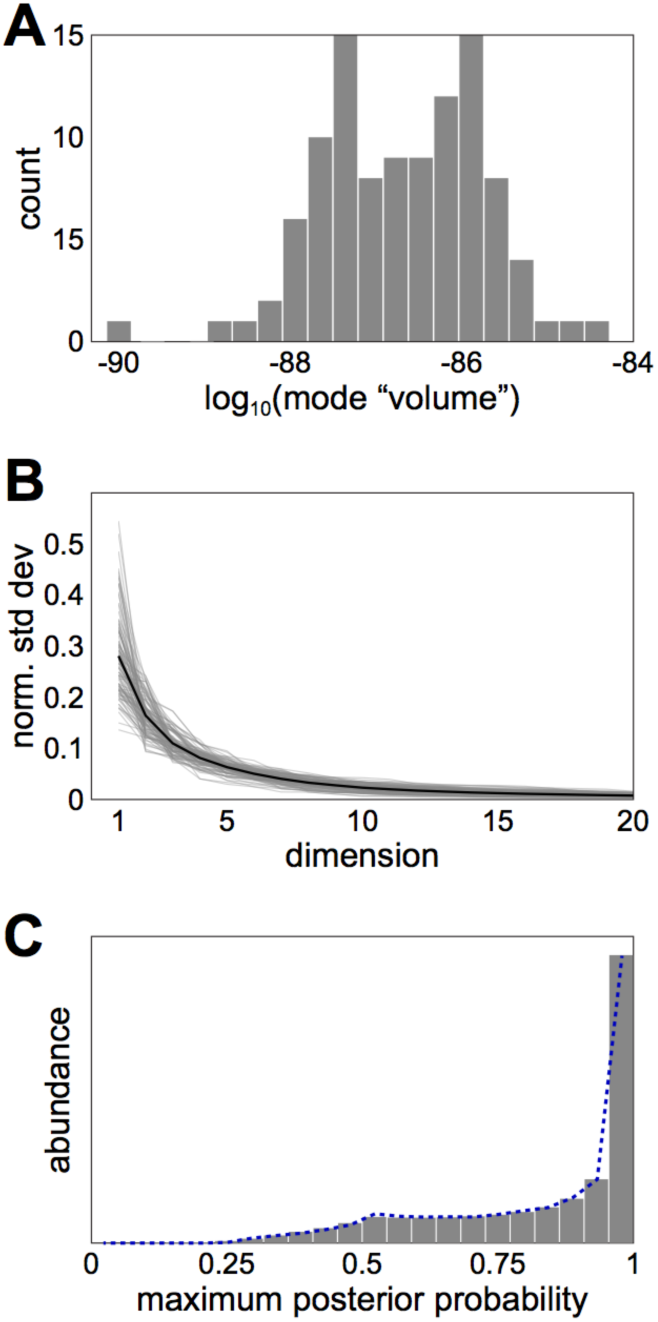
Statistical characteristics of clusters produced by PCA_20_-GMM –. **A**)Histogram of mixture component “volumes” for the *k* = 104 clustering of the data from fly experiment 37_1_. Volume computed as the product of the eigenvalues of each Gaussian mixture component covariance matrix. **B**) Relative size of each of the dimensions of the 104 mixture components of fly experiment 37_1_. For each component, the eigenvalues of the covariance matrix were sorted, normalized by their total, and plotted. Grey lines are specific components, the black line is their average. **C**) Histogram of the maximum posterior probability of all mixture components across all frames of all fly experiments. The posterior probability of component *a* at point *x* is defined as the probability density of component *a* at *x* divided by the sum of the densities of all components at *x*. Dotted blue line is the equivalent histogram of maximum posterior probabilities achieved by the arrangement of mixture components in Figure 5A.

To visualize the overlap between our clusters, we created a set of toy data and arranged it (via a Monte Carlo method) to have the same distribution of posterior probabilities Todd, Kain & de Bivort, 2016 – preprint version as our experimental data (Figure 5A). Based on this we believe our clusters are in general relatively far apart from each other, but that some are not Gaussian in shape. GMM may therefore need to invoke several mixture components in order to fit what is a single (non-Gaussian) cluster (Figure 5B). A watershed transform can identify non-Gaussian modes more naturally, but due to its memory requirements, we weren’t able to run it in our 20-dimensional compressed postural dynamics space. As an approximation of a high-dimensional watershed transform, we applied what we termed a *sparse watershed* algorithm, which maps each data point to a local maximum by ascending the gradient (Snyman 2005) of the GMM distribution. Each data point (assigned to a *child cluster* by PCA_20_-GMM) was then reassigned to a *parent cluster* by taking the maximum posterior probability at its local maximum. To further ease the computational requirements, we ran this procedure on only the cluster means and assigned all points in each cluster to the same parent cluster, which yielded approximately the same result as transforming each point separately (Figure 5C).

**Figure 5.**
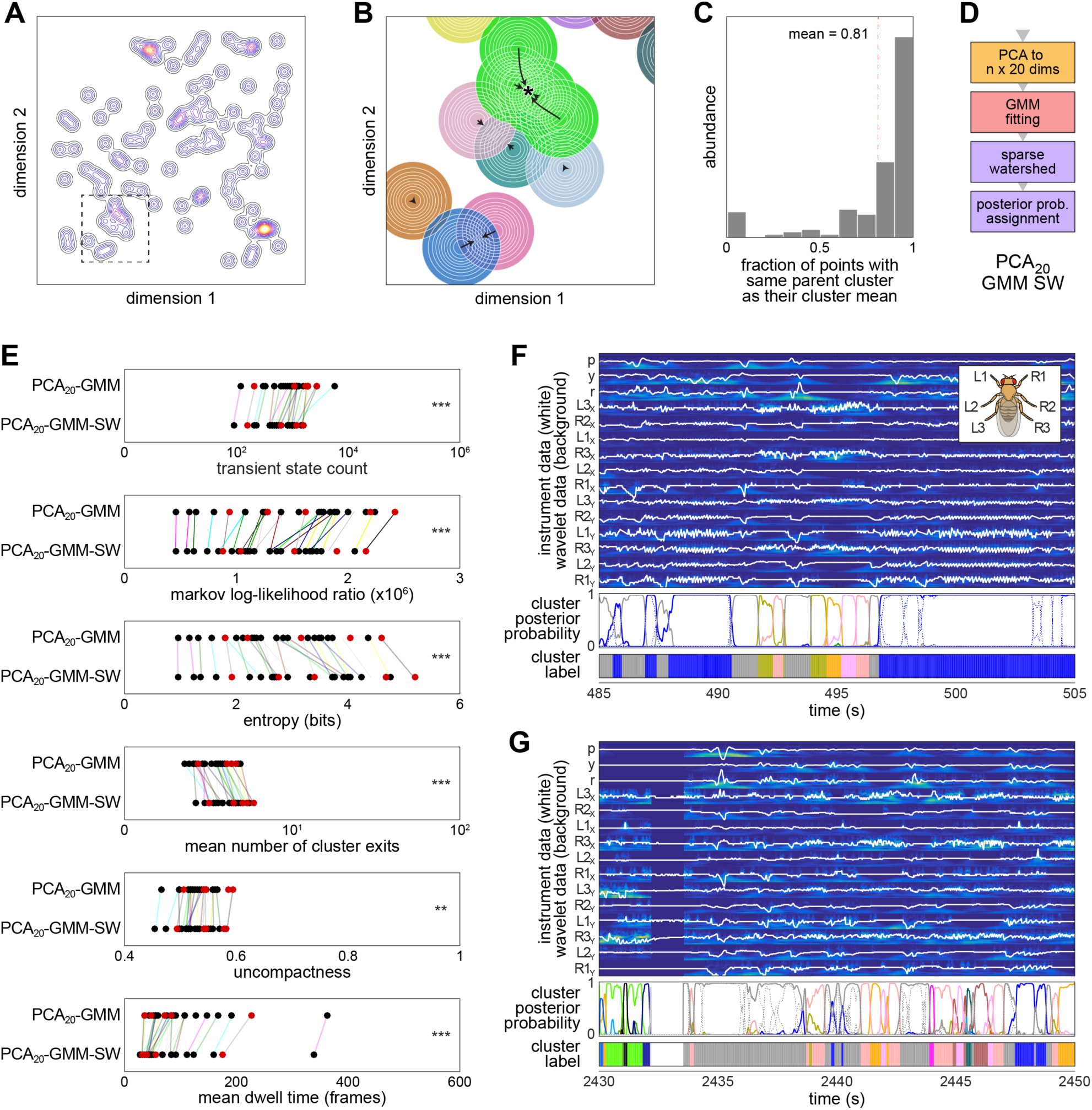
Use of sparse watershedding to consolidate Gaussian mixture components –. **A**) Contour plot of a 2D Gaussian mixture model with 100 components, all of which have identical weights and the identity matrix as a covariance matrix. This arrangement of components has essentially the same distribution of maximum posterior probabilities as produced by the PCA_20_-GMM method on real fly data (Figure 4C). This toy example serves to visualize the sparseness of the mixture components that produces the observed distribution of maximum posterior probabilities. Inset expanded in B). **B**) lllustration of how the sparse watershed algorithm ascends the Gaussian mixture probability density to local maxima (arrows). Green components whose means (origins of arrows) ascend to the same point (asterisk) are consolidated. The sum of the separate components shown here is the inset of A). **C**) Histogram showing percentage of data points sampled from each mixture component for which sparse watershedding yields the same result as for the cluster mean. Dashed red line indicates the mean across all components (81%). **D**) Extended mapping method flow chart characterizing the PCA_20_-GMM-SW mapping method. **E**) Metric values for all fly experiments using PCA_20_-GMM and PCA_20_-GMM-SW. Asterisks indicate statistical significance as assessed by Wilcoxon signed-rank tests and corrected for 6 comparisons: *** *p*<0.001; ** *p*<0.01. **F**, **G**) Examples of 20 s frame sequences from fly experiment 37_1_ illustrating the data values after pre-processing (white lines) superimposed on their wavelet time-frequency representation. p = pitch, y = yaw, r = roll. L1X indicates the x-component of the position of the left (L) fore leg (1). Inset diagram shows leg labels on a fly in top-down view. Middle panel is the posterior probability associated with each frame of each PCA_20_-GMM-SW cluster. Parent cluster posterior probabilities (solids) are the sum of their respective child cluster posterior probabilities (e.g. dotted blue lines sum to the solid blue line). Bottom panel is the cluster label applied to each frame by PCA_20_-GMM-SW.

The sparse watershed algorithm allows us to consolidate GMM-assigned clusters (c.f. several alternative ways to do this depending on the data set; Hennig 2010, Garcia *et al* 2010), leading to a refined mapping method which we refer to as PCA_20_-GMM-SW (Figure 5D). We recomputed our metrics by running the PCA_20_-GMM mapping method with cluster counts chosen to match the PCA_20_-GMM-SW consolidated cluster counts (Figure 5E). We found that every metric was significantly different between PCA_20_-GMM and PCA_20_-GMM-SW. In half of the metrics, the addition of sparse watershedding shifted the metric values in the favorable direction: transient state count (decrease), entropy (increase), and uncompactness (decrease). The other metrics changed in the unfavorable direction: markov log likelihood (decrease), mean number of cluster exits (increase), and mean dwell time (decrease).

On balance, based on metrics alone, there is no definitive advantage to using sparse watershedding. However, there are cases (as in Figure 5B) where sparse watershedding can consolidate modes which are invoked by GMM to increase the quality of fit to a particular peak of behavioral probability density. In the absence of a consolidating step like sparse watershedding, these “shaping” modes are undesirable since they do not characterize new behavioral modes. Thus, we conclude that the flexibility of watershed-style algorithms to identify the boundaries of local maxima in behavioral probability density non-parametrically is an advantage. The ambivalent aggregate difference in metrics between PCA_20_-GMM and PCA_20_-GMM-SW mean we can freely invoke this procedure, gaining its flexibility at no obvious cost to the overall quality of the unsupervised clustering. Several examples of sparse watershedding working to consolidate similar clusters can be seen in Figure 5F, G.

Once frames are assigned to clusters via PCA_20_-GMM-SW we can construct a Markov transition matrix to explore the dynamics within behavioral space. After sorting the rows and columns of the transition matrix of fly experiment 37_1_ by similarity, several structural aspects emerge (Figure 6A). The most evident is the partitioning of the space into two subdivisions of roughly equal size. The distribution of energy over dimensions for each cluster (Figure 6B) allows us to recognize the upper rows as containing motion in both front and rear legs (e.g. wing grooming), while the lower rows contain energy in only the front legs (e.g. front leg grooming). The three center rows contain relatively populous clusters (Figure 6C): one low-energy cluster (Figure 6A-C, red tabs) and two with energy in all legs, reflecting running (one of which is denoted in Figure 6A-C, green tabs). The transition matrix shows which sequences of behavior are possible. For example, flies evidently do not transition directly from running to front leg grooming, but instead transition through an intermediate state (e.g. the resting state).

**Figure 6.**
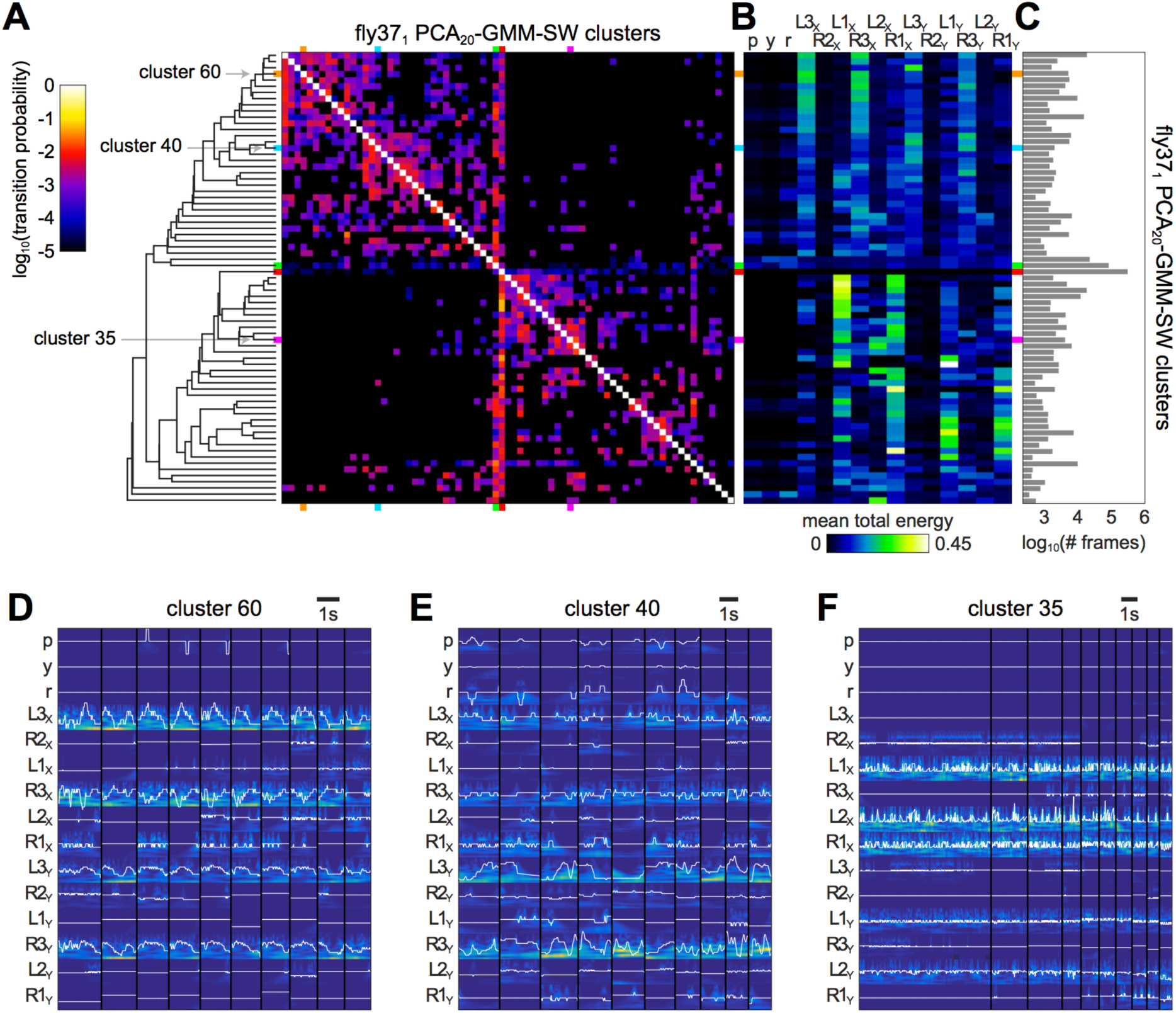
Clustering of behavioral space for fly experiment 37_1_ by PCA_20_-GMM-SW –. **A**) Markov transition matrix between the inferred clusters of fly experiment 37_1_. The entry in the *i*^th^ row, *j*^th^ column is the probability that a frame in cluster *i* will be followed by a frame in cluster *j*. Rows and columns have been arranged by similarity as indicated by dendrogram on the left. The rows and columns corresponding to clusters 60, 40 and 35 are marked with colored tabs, as are the clusters corresponding to rejected low-variance frames (red) and a cluster representing running behavior (green). **B**) Heatmap of the mean total wavelet energy of each input dimension (columns) for those frames receiving each cluster label (rows). Data vector labels as in Figure 5F, G. **C**) Abundance of each cluster label in fly experiment 37_1_. **D**-**F**) Examples of streaks of consecutive frames receiving cluster labels 60, 40 and 35 respectively. Cluster 60 labels frames with motion predominantly in the x-coordinates of the hind legs. Cluster 40 labels frames with comparatively disorganized motion predominantly in the y-coordinates of the hind legs. Cluster 35 labels frames with high frequency motion predominantly in the fore legs and middle leg on the left. Vertical black lines demarcate distinct frame streaks.

To demonstrate the similarity of behaviors assigned to the same cluster and to examine the data more closely, streaks of frames labeled with each of three clusters are shown in Figure 6D-F. The distribution of lengths of streaks of consecutive frames receiving cluster label 35 appears to be roughly exponential. Indeed, this appears to be true for all clusters (data not shown). Thus, the process of leaving behavioral modes may be approximately memoryless (Murphy *et al* 2002).

To examine the evolution of behavior over time we plotted a 10 s sequence of frames in behavioral space, visualized with the corresponding tSNE2-watershed density map for reference in the background (Figure 7). The clusters identified by PCA_20_-GMM-SW are spread across multiple tSNE2-watershed regions, illustrating the extent of discordance between these methods. As others have found (Berman *et al* 2014, Crall *et al* 2016), the step size distribution appears to be bimodal: the animal makes generally small steps within a behavioral cluster and large steps in moving between them.

**Figure 7.**
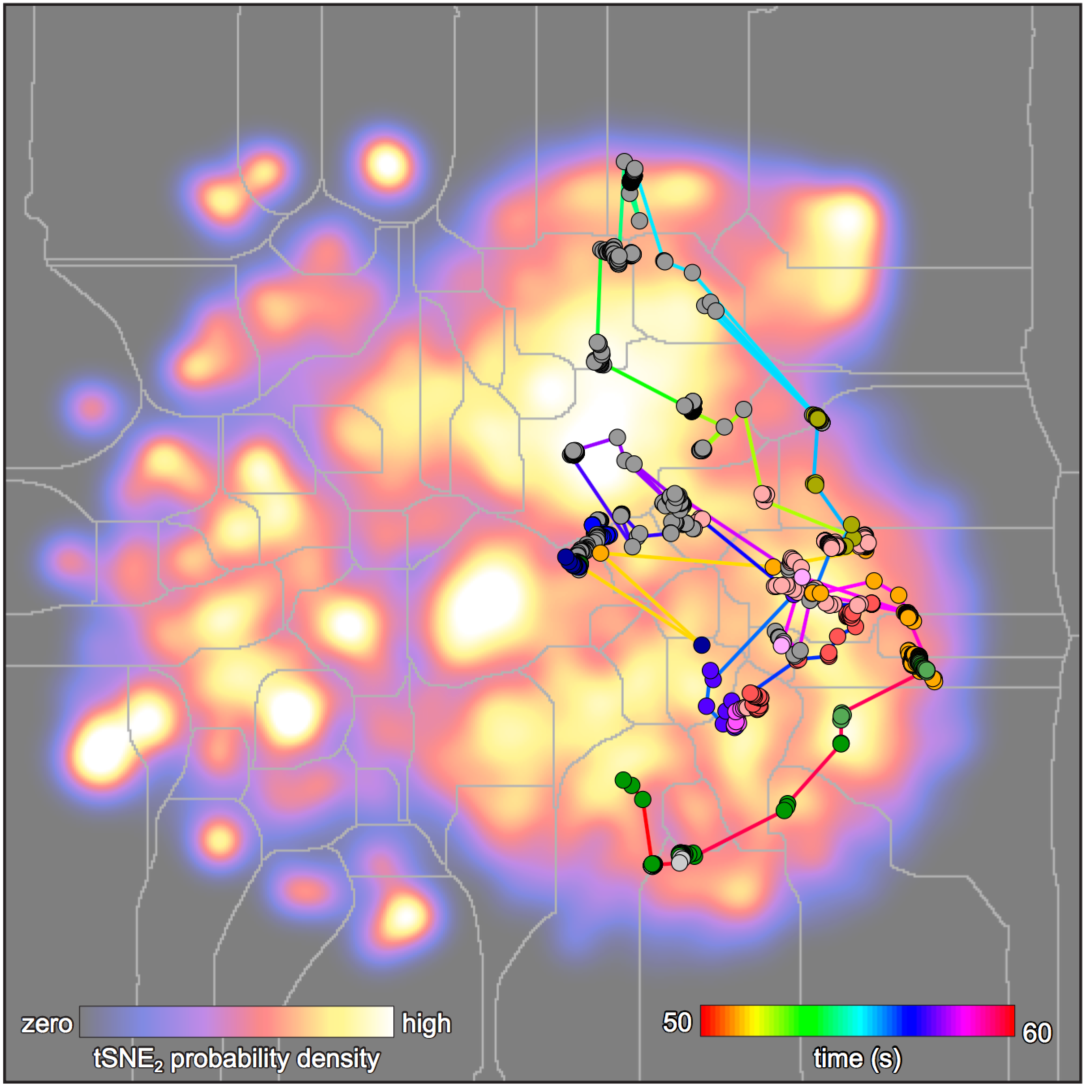
Time evolution of 10 s of data from fly experiment 37_1_ –. Background heatmap is a density estimate of points embedded in two dimensions using t-SNE. Grey lines are watershed boundaries. Thus the regions demarcated in this heatmap correspond to the cluster boundaries of the tSNE2-watershed method (Berman *et al* 2014). Line segments of the foreground data are time-coded. Foreground points are spatially positioned by the t-SNE embedding, but have markers whose color indicates the cluster label assigned to those points by the PCA_20_-GMM-SW mapping method.

### Inter-fly comparisons

Having established the effectiveness of PCA_20_-GMM-SW on a single fly, we proceeded to apply it to all flies simultaneously. By fitting the GMM on data sampled from all flies (“co-fit”), we used PCA_20_-GMM-SW to establish a behavioral space for all of our data taken together. To determine the value of *k* used in fitting the GMM to this aggregated data set, we used PCA_20_-GMM on the data from each fly independently. We systematically varied *k* in each case and computed the Bayesian Information Criterion (BIC) of the GMM fit (Schwarz 1978). This assesses the likelihood of the data under the GMM, assessing a penalty for the number of fitting parameters. Knees in the curve of BIC vs *k* can indicate when the addition of Gaussian mixture components offers no (or only marginal) fit improvement (Salvador and Chan 2004). We found that across flies, a soft knee was present around *k* = 70 mixture components. Therefore we sought to produce an aggregate clustering with approximately 70 labels in it. We found that PCA_20_-GMM-SW consolidated mixture components more aggressively in the co-fit data set than it did in individual fly data sets. In the end, we ended up choosing a PCA_20_-GMM-SW clustering in which 180 GMM components consolidated to 40 clusters (plus the additional low-variance cluster).

Many of the clusters defined by this mapping method correspond to well-known behavioral elements in flies, such as a specific kind of fore leg grooming (cluster #15; see Supplemental Movies M2 and M3). However, some behavioral clusters captured behavioral sequences that seem like the concatenation of two distinct behaviors. For example, cluster #7 appears to represent a rotation of the floating ball followed by hind leg grooming. This pattern appears similarly in all the flies examined (see Supplemental Movies M2 and M4). Thus, the unsupervised approach appears to be identifying behavioral patterns that are distinct from those a human investigator might define.

We examined cluster size distributions across fly experiments (Figure 8A) sorting rows (flies) and columns (clusters) by their respective similarity. This reveals a broad consistency between flies in the overall distribution of time spent in each cluster (row). There are also apparent differences between individual flies, since multiple trials from the same fly tend to be grouped together. The total number of frames labeled by each cluster varied substantially across clusters, indicating that PCA_20_-GMM-SW is flexible enough to identify both rare and common modes of behavior. The distribution of wavelet energy over raw data dimensions across our 40 co-fit clusters (Figure 8B) is largely concordant with the distribution across 73 clusters for fly experiment 37_1_ (Figure 6B). Interestingly, sorting cluster energy distributions by the same permutation that sorts clusters by similarity in their abundance across flies (Figure 8A) moves together clusters which have similar distributions of wavelet energy. The fact that variation in cluster abundance across individuals (Figure 8A) is somewhat concordant with cluster variation in wavelet energy (Figure 8B) implies that there may be common mechanisms determining 1) which legs move in a cluster and 2) variation across individuals.

**Figure 8.**
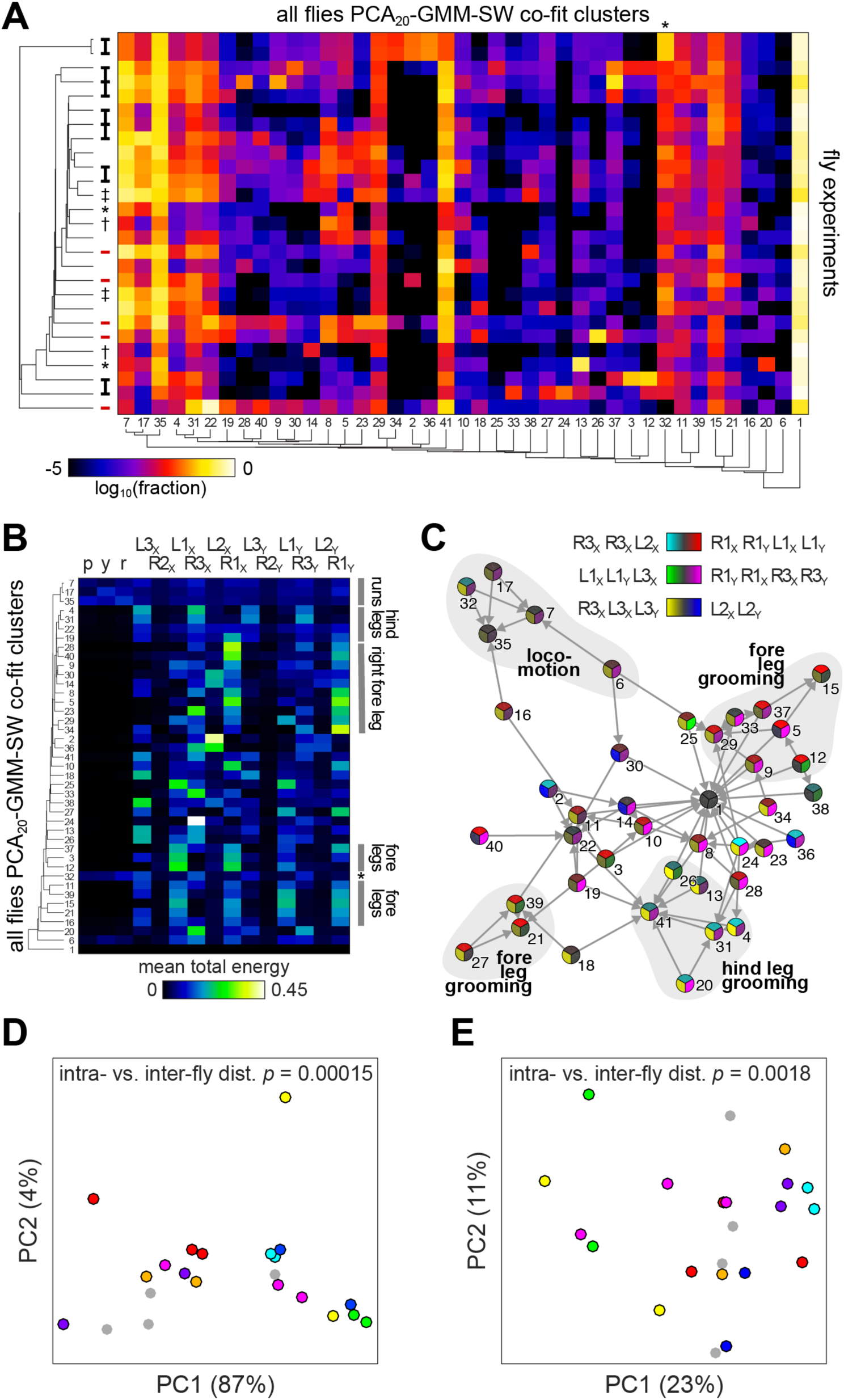
Comparison across fly experiments using a common PCA_20_-GMM-SW mapping –. **A**) Heatmap of the fraction of frames from each fly experiment (rows) receiving each cluster label in the PCA_20_-GMM-SW co-fit mapping. Rows and columns are respectively sorted by similarity as indicated by the left and bottom dendrograms. Bold black brackets next to the row dendrogram indicate experiments on successive days from the same fly which were placed adjacently by the dendrogram, i.e. having similar profiles of cluster labels. Dendrogram tips with *, † and ‡ represent experiments from the same fly that were not placed adjacently. Red tabs indicate *nan* mutant flies. Numbers at bottom indicate presorting cluster label numbers. Asterisk indicates the one behavioral cluster that was significantly different in abundance (*p* = 0.013) in *nan* animals compared to wild type. **B**) Heatmap of the mean total wavelet energy of each input dimension (columns), for those frames receiving each cluster label (rows) from the PCA_20_-GMM-SW co-fit mapping. Rows sorted by similarity as indicated by dendrogram, numbers and asterisk as in A). Grey bars at right indicate blocks of clusters which have similarly distributed wavelet energy, as determined by inspection. **C**) Directed graph of the transitions between cluster labels in the PCA_20_-GMM-SW co-fit mapping. All frame transitions for all fly experiments were pooled here. Node positions were determined by the force-directed algorithm (Kobourov 2012). Each node is divided into three wedges colored to reflect the first three principal components of the mean total wavelet energy across input dimensions (i.e. the first three PCs of panel B). Cyan vs. red coloration in the top wedge indicates greater energy in hind legs vs. fore legs (PC1, 36% of variance). Green vs. Magenta coloration in the bottom-right wedge indicates greater energy in the left vs. right legs (PC2, 18%). Yellow vs. blue coloration in the bottom-left wedge indicates greater energy in the middle left leg vs. the hind legs (PC3, 15%). In the key, input dimensions are listed if they received PCA loading values with absolute values > 0.2. Color scales are offset so that their values in the low-variance frames are neutral grey. Directed edges indicate frame-to-frame transitions more probable than 10^−1.8^. Grey regions in the background are high-level cluster categories determined by inspection. Numbers as in A) and B). **D**) Plot of the first two PCs of the cluster label distribution (i.e. panel A) for each fly experiment. Points with the same color are sequential experiments from the same fly. P-value is the statistical significance of the difference in the mean distance (in the original 41 dimensional cluster label space) between pairs of experiments either within fly or between random pairs of experiments, tested using the Wilcoxon–Mann–Whitney test. **E**) As in D), but for data consisting of the 1681 Markov transition probabilities between cluster labels for each fly experiment.

We constructed first-order Markov transition matrices for each fly in the co-fit behavioral space, then we pooled them together to explore co-fit behavioral dynamics. To visualize the connections between behavioral states we laid them out in a force-directed graph with node edges representing the most prevalent transitions (Figure 8C). A general structure emerges with a central hub consisting of the low-variance rest state (cluster #1) and several partially connected outlying subnetworks of nodes suggesting a hierarchical organization of behaviors. Nodes representing behaviors involving the same combinations of leg movements (similar color patterns) are more likely to link to one another, consistent with the transitions for fly experiment 37_1_ shown in Figure 6A, B. This visualization also makes it apparent that there may be a bias toward energy on the right side of the animal (more magenta than green wedges in Figure 8C). This is also discernible in the wavelet energy distributions of each cluster (Figure 8B), possibly reflecting population-level behavioral handedness.

We also projected individual fly experiments into a two-dimensional space consisting of the first two principal components of individual 41x1 cluster abundance distributions (Figure 8D) and individual 41x41 transition matrices (Figure 8E). This shows a statistically significant grouping of experiments from individual flies, consistent with the fly individuality observed above in Figure 8A.

Lastly, we examined *nan* mutant animals vs wild type. Consistent with subjective observations made during experiments, we found that a cluster representing a locomotor behavior (#32) was statistically significantly (*p* = 0.013) less abundant in *nan* animals than in wild type. Surprisingly, this was the only behavioral cluster that differed in frequency between the genotypes, after correcting for multiple comparisons.

## DISCUSSION

We set out to compare alternative approaches to unsupervised behavioral mapping and apply the most promising method to our leg-tracker data from wild type and *nan* animals. Selecting among competing unsupervised mapping methods on the basis of performance metrics seems paradoxical, since this will introduce bias in the outcome of the unsupervised mapping procedure, which, by definition, attempts unbiased clustering. However, we believe that the criteria we used to evaluate alternative mapping methods (our metrics) are general enough so as not to adversely distort the output of the mapping method comparisons. We also tried to be conservative in assigning normative value to the metrics, e.g. we said that high entropy classifications (i.e. those that distribute frame assignments more evenly across clusters) are *probably* desirable. We found that no single mapping method performed best across all these metrics. In fact, three principle components are needed to explain 95% of the variance in mapping method rank across all the metrics in Figure 3. So, there is likely no single method among the six we examined here that can be applied without compromises. Moreover, the suitability of mapping methods may be data set dependent, a possibility we did not explore here.

The PCA_20_-GMM method performs well in all metrics except the mean number of cluster exits. This metric, which we assume to be probably desirable, characterizes the average number of clusters that follow any particular cluster in the sequence of labeled frames. We assume that fewer such exits is better because this makes the entire sequence of labels more predictable. PCA_20_-GMM was ranked fifth of sixth in this respect; in all other metrics it was ranked first or second.

Because there is no particular reason to think that behaviors are manifested as multivariate Gaussians in real life, we implemented the sparse watershedding extension so that GMM components contributing to a single mode of density would be consolidated into a single cluster. To compute this consolidation we identified those mixture components whose means flowed (in the watershed analogy) to the same points in 20-dimensional space using a gradient ascent algorithm. This is not a trivial calculation, requiring the calculation of a 20-dimensional PDF at each step of the gradient ascent, for each of ∼100 mixture component means, for each of the 27 fly experiments. Only computing the sparse watershed for the component means is a compromise under these computational constraints, as it could, in principle, be performed for every point in the data set. That said, our analysis of the sparse watershed behavior of points randomly sampled out of the mixture components of fly 37_1_ (Figure 5C) suggests that a large majority of points drawn from a mixture component will consolidate to the same cluster as their component mean. This is consistent with the relatively low-density packing of mixture components in our mapping space (Figures 4C, 5A).

The degree of consolidation observed by the SW algorithm diverged between the individual fly data sets and the co-fit aggregated data. In the case of the latter, the fold-reduction in number of clusters was much higher, and appeared to scale sub-linearly with the *k* used for the initial GMM step (data not shown). Perhaps this reflects a case where the superimposed modes from each fly were slightly misaligned, meaning that more initial mixture components were needed to fit them, but they were still close enough to consolidate together via sparse watershedding. Alternatively, combining data across individuals may average away differences (Figure 8D, E) to create a species-level data set that behaves differently under these procedures.

The PCA_20_-GMM-SW method is by no means the most sophisticated possible approach. For example Wiltschko *et al*. (2015) fit a hidden Markov model directly to the data instead of inferring it after clustering as we have done. Thus, their optimization is simultaneously on the Markovian transitions between states as well as the shape of the states themselves. It also produces a generative model which can synthesize new data sets instead of simply analyzing existing data. Our approach prioritizes the mapping of the total space occupied by postural dynamics rather than their transitions. The assumption of Markovian transitions between states is in general reasonable, and we found that the distribution of lengths of streaks of frames receiving the same cluster label were roughly exponential, as expected from a memoryless Markov process. However, there are biological reasons to think that behavioral transitions may not be Markovian, namely state-dependence and neuromodulation, which act on timescales longer than typical behavioral transitions, but shorter than our experimental recordings (e.g. Cohn *et al* 2015).

One central finding is that clusterings computed in higher dimensions appear to perform better in our metrics than clusterings computed in two dimensions. This may simply reflect the inevitable compression or distortion of high-dimensional data when it is embedded in two dimensions. High dimensionality poses many challenges, among them computational complexity and the inability to confirm intuition by examining the data visually. The examination of metrics that simply characterize mapping output without much normative weight (e.g. whether clusters are uncompact or have long vs. short dwell times) allowed us to build intuition about the space of behaviors in higher dimensions than can be visualized. Likewise, using a Monte Carlo method to arrange Gaussian mixture components in two dimensions to have the same statistical distribution of maximum posterior probabilities as in 20 dimensions (Figures 4C, 5A) also allowed us some intuitive access to the high-dimensional representation of our data set.

Based on these experiences working with our data set, we recommend the method outlined in Figure 9 for behavioral clustering. If the computational resources are available, GMM-BIC curves provide statistical support for the choice of mixture component count prior to the sparse watershedding step (i.e. looking for a knee in the BIC-vs-*k* curve). As an alternative, the tSNE_2_-watershed pipeline allows the mapping output to be visualized, which may provide a basis for choosing *k* for subsequent analysis. For example, one might find that there is a range of 2D Gaussian blurring kernel radii which give very similar numbers of watershed clusters. This would be a sign that the map is robust to this free parameter in the density estimation step. However, for our data set we found that the number of watershed clusters was roughly linearly proportional to the radius of the blurring kernel, so we could not use tSNE2-watershed to build a strong prior on the number of clusters in the data.

**Figure 9.**
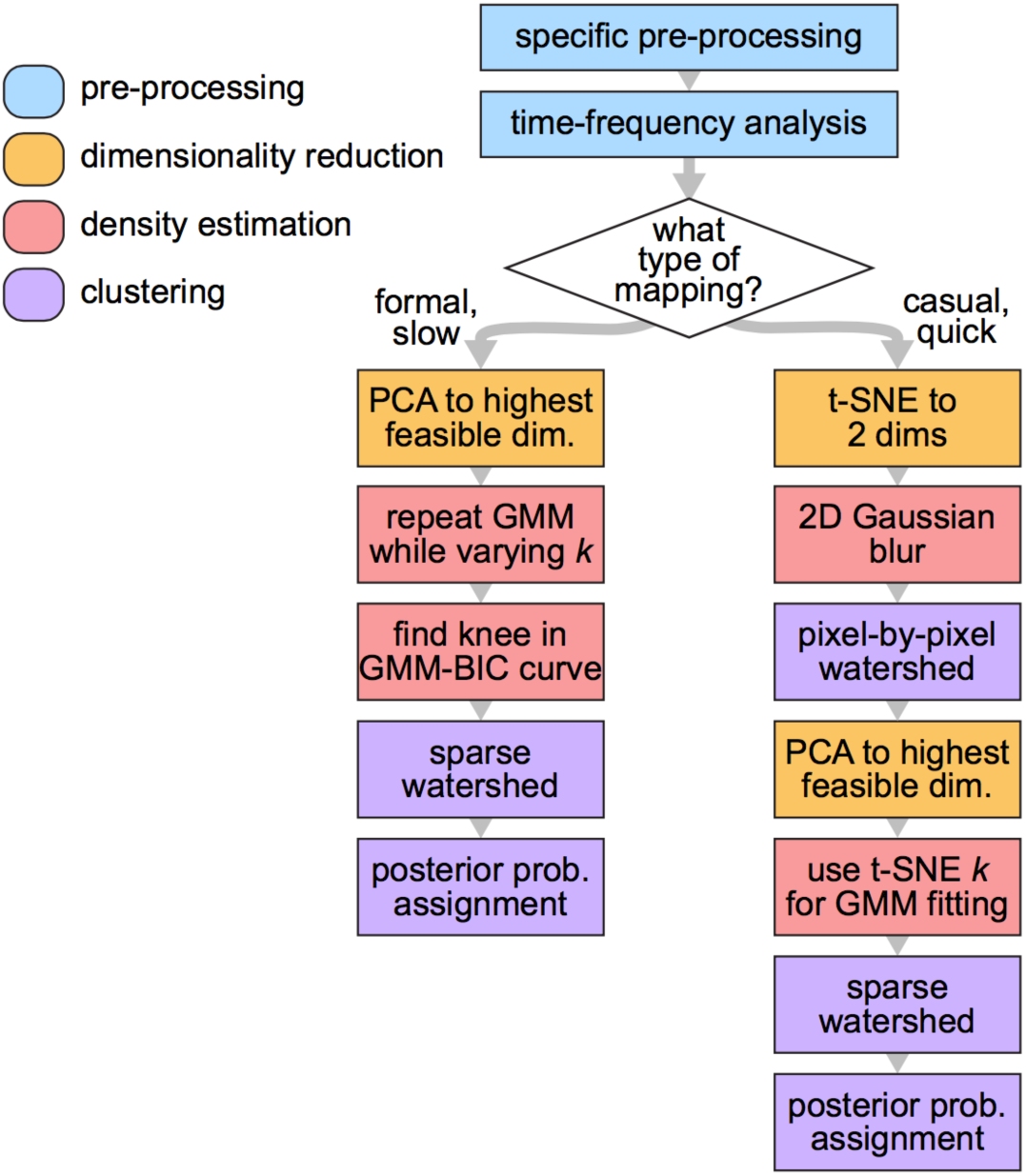
Suggested unsupervised behavioral mapping method flow chart –. The right branch represents an option for computing fast clustering by PCA-GMM-SW using a value of *k* informed by a first round of exploratory analysis using tSNE_2_-watershed. By contrast, a formal, methodical approach (left branch) stipulates systematically varying the value of *k* used in GMM and searching for a knee in the GMM-BIC curve to determine a final PCA-GMM-SW mapping *k*.

Combining data from all the fly experiments allowed us to make a common clustering for comparison across flies, similar to Berman’s method (2014) for co-embedding multiple animals’ data in a single mapping. This allowed us to compare wild type and *nan* mutant animals. Surprisingly, we only found a single behavioral cluster that had a significantly different abundance between these genotypes. Cluster 32, which encodes a behavior that features both hind leg grooming and walking, was reduced in *nan* flies. Such a difference was predicted at the time of experiments because the *nan* flies seemed clearly more lethargic. In fact, the small sample size of five *nan* recordings was due largely to very few of these animals performing on the floating ball. It is a surprise to see that outside of this one cluster, the abundance of behavioral clusters in *nan* animals is generally indistinguishable from that of wild type. This suggests that *nanchung*, and perhaps proprioception, may be dispensable for the establishment of full repertoires of behavioral building blocks, despite the necessity of this gene for the execution of locomotor behaviors (Mendes *et al* 2013, Isakov *et al* 2016).

With the co-fit PCA_20_-GMM-SW mapping we also found evidence of largely similar behavioral cluster abundances across individuals, but at the same time, significantly greater similarity across trials within a fly than between flies. The pairwise transition rates between clusters appeared to be idiosyncratic as well. These findings are consistent with behavioral individuality in *Drosophila* phototaxis (Kain *et al* 2012), locomotor handedness (Buchanan *et al* 2015) and thermal preference (Kain *et al* 2015). However, the clear evidence of individuality in PCA_20_-GMM-SW cluster abundance distributions is notable when compared to the lack of signal of such individuality in the supervised classification results of Kain *et al* (2013). This indicates that despite the challenges in evaluating and implementing unsupervised methods, relatively unbiased clustering approaches may indeed live up to their promise of providing more power to map behavior.

## CONCLUSION

In this work we conducted the first systematic exploration of alternative unsupervised methods for mapping the behavior of *Drosophila*. We devised seven different metrics that range from definitely desirable to possibly desirable for the output of an unsupervised clustering method. Using these we were able to compare the overall performance of seven combinatorial alternative mapping methods. We concluded that keeping the data in as many dimensions as possible for clustering is preferable, at least for our data set. Implementing this method on merged data from multiple individuals and genotypes, we saw that while individual flies implement very similar motor repertoires, there are repeatable differences in the abundances with which they implement behavioral clusters as well as the transitions they use to generate higher order sequences of behaviors. Surprisingly there was little difference between the repertoires of wild type and proprioceptive mutant flies. Future work building on these results will include: 1) comparison of the methods we considered here with altogether different approaches, such as structure learning (Vogelstein *et al* 2014), 2) characterization of the generality of these findings beyond data from the leg-tracking instrument, especially video data, 3) improvement of our PCA_20_-GMM-SW method, e.g. by the implementation of the sparse watershedding algorithm at all data points, and 4) sensitivity analysis of PCA_20_-GMM-SW under variation in noise in the data, dimensionality, sample sizes, or other factors. This sensitivity analysis could be conducted using hidden Markov models (Wiltschko *et al* 2015) to generate plausible sequences of realistic but synthetic data whose noise, dimensionality and sample size parameters can be adjusted arbitrarily.

## ACKNOWLEDGEMENTS

Thanks to Mike Burns for being a helpful sounding board for early ideas. We owe Eric Stone a skyr of thanks for his help in thinking through the formulations of candidate metrics. Zach Werkhoven provided thoughtful comments on the manuscript, kirimvos. We are grateful to the Elizabeth Kane Lab for supporting the collaboration on this project. We acknowledge the Rowland Junior Fellows Program and the Alfred P. Sloan Foundation for generous funding support.

